# Bayesian model discrimination for partially-observed epidemic models

**DOI:** 10.1101/646067

**Authors:** James N. Walker, Andrew J. Black, Joshua V. Ross

**Affiliations:** School of Mathematical Sciences, University of Adelaide, Adelaide, SA 5005, Australia; ACEMS, School of Mathematical Sciences, University of Adelaide, Adelaide, SA 5005, Australia

**Keywords:** Model selection, Epidemic modelling, Importance sampling, Markov chain, Particle filter

## Abstract

An efficient method for Bayesian model selection is presented for a broad class of continuous-time Markov chain models and is subsequently applied to two important problems in epidemiology. The first problem is to identify the shape of the infectious period distribution; the second problem is to determine whether individuals display symptoms before, at the same time, or after they become infectious. In both cases we show that the correct model can be identified, in the majority of cases, from symptom onset data generated from multiple outbreaks in small populations. The method works by evaluating the likelihood using a particle filter that incorporates a novel importance sampling algorithm designed for partially-observed continuous-time Markov chains. This is combined with another importance sampling method to unbiasedly estimate the model evidence. These come with estimates of precision, which allow for stopping criterion to be employed. Our method is general and can be applied to a wide range of model selection problems in biological and epidemiological systems with intractable likelihood functions.

## 1. Introduction

Biological and epidemiological processes can have highly complex behaviour and in many cases we do not observe all of their important features. As our observations may depend on features we do not see, it can be difficult to infer properties of this partially-observed process. To add complexity to the problem, there may be several theories about the behaviour of the process and it may not be obvious as to which of these models is most appropriate given some set of observations. For example in epidemiology, it is widely recognised that the shape of the infectious period distribution is critical to understanding historical disease incidence, and also for accurate evaluation of control measures for public health use [1, 2]. Similarly, it has been identified that the relative timing of symptoms and infectiousness is a key determinant of ability to control an outbreak [3]. Investigating these issues is most easily done by encoding certain features in the structure of the model itself rather than just parametrising it, hence to discriminate different possibilities we need to perform some type of model selection.

Bayesian model selection compares models using the probability that each model generated the data given some observations and prior distribution over models and parameters [4]. We consider the model to be inferred as a parameter (typically with a uniform prior) and hence multiplying the model evidences, that is, the likelihood of each model, by the prior distribution and normalising gives the probability of each model given the data [5]. While the interpretability and the consistency with the Bayesian paradigm is desirable, Bayesian model selection has some issues: a large amount of data may be required before models can be distinguished; and the evidence is typically difficult to calculate as the likelihood is intractable. This is true for epidemic models, which for small populations are most naturally represented as partially-observed continuous-time Markov chains (CTMCs) [6, 7, 8].

In this paper we describe an efficient way of performing Bayesian model selection for partially-observed CTMCs and apply our method to the two problems in mathematical epidemiology discussed above: inferring the shape of the infectious period distribution and identifying the onset of symptoms relative to infectiousness. Since the 2009 outbreak of swine flu’, countries around the world, such as the USA, UK and Australia, have outlined data collection protocols for pandemic infectious diseases [9, 10, 11]. The datasets considered in this paper are of the form that will be obtained during such an outbreak in Australia, as outlined in the Australian Health Management Plan for Pandemic Influenza [11]. These datasets consist of daily counts of symptom onset stratified by household. A study of this kind has not yet been enacted, so we conduct simulation studies; this also allows us to validate results. Our method works by calculating unbiased estimates of the likelihood via sequential importance resampling, which in turn is used in another importance sampling algorithm to estimate posterior model probabilities. A novel feature of this method is that the likelihood is estimated via a scheme which is ideally suited for partially-observed CTMCs, where one component of the state is observed exactly [8]. This works by sampling realisations of the partially-observed process in a way that realisations always match with observations. Our combined method is both computationally efficient and embarrassingly parallelisable and hence well suited to implementation on modern computing hardware. Further, the ability to estimate error bounds and use stopping criterion ensures the accuracy of evidence estimates. Similar CTMC models see wide use in areas of biology such as phylogenetics [12], ecology [13] and cell biology [14], and hence our method may be applied to a broad range of biological model selection problems. Matlab code is provided for all algorithms developed in this paper as part of the EpiStruct project [15].

We first discuss the methods for importance sampling over the parameter space, and the sequential importance resampling algorithm used to estimate the likelihood in Section 2. We then describe the two case studies, their implementation and results in Sections 3 and 4. Lastly we conclude in Section 5 and discuss how our method relates to those already in the literature.

## 2. Methods

Importance sampling is a method for estimating properties of one distribution by sampling from another distribution; the bias is corrected to obtain unbiased estimates related to the distribution of interest [16]. This technique is useful when the target dis-tribution is difficult to sample from, such as the distribution of latent variables from a partially-observed Markov chain. This method has been effectively applied for parameter inference, for example in particle marginal MCMC schemes [17, 18] and has also been applied to model selection [4, 19, 20, 21]. In this paper we use importance sampling in the space of latent variables to estimate likelihoods via sequential importance resampling and within the parameter space to calculate the model evidence. The novelty of this approach is that the likelihood is estimated via the most suitable kind of importance sampling scheme for these kinds of models, as described in [8].

Here we introduce importance sampling for model selection, detail how likelihood estimates are used in model selection and describe how sequential importance resampling is used for likelihood estimation.

### 2.1. Importance sampling for evidence estimation

Importance sampling for model selection has been discussed in [4, 19, 20], and in this paper we adopt a similar approach. However, our implementation uses an efficient particle filter to obtain unbiased estimates of the likelihood, which allows us to obtain unbiased estimates of the evidence.

Suppose we have data set *y* of observations from a model with parameter set *θ* in parameter space Θ. Let *p*(*θ*) denote the prior distribution and *p*(*y|θ*) denote the likelihood function, then the model evidence is given by

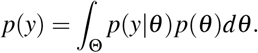

To estimate *p*(*y*) we may sample *m* parameter sets *θ*_1_*, …, θ_m_* from some arbitrary, importance sampling, density, *q*(*·|y*), with support Ψ where Θ ⊆ Ψ. We then compute random variables

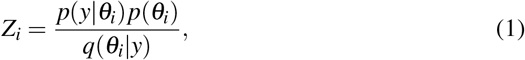

for *i* = 1*, … m*, where *p*(*θ*) is the prior distribution and *p*(*y|θ*) is the likelihood function. These random variables are chosen so that their expectation is given by

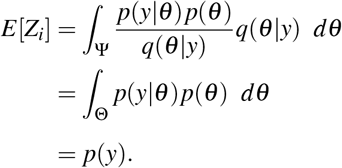

Hence, the mean of the *Z_i_*’s provides an unbiased estimate of the evidence, 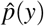. If the likelihood is intractable we can use unbiased estimates, 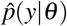, instead of *p*(*y|θ*) in Equation (1), by the law of total expectation. Note that using importance sampling to estimate *p*(*y*) involves obtaining independent identically distributed samples of *Z_i_* and taking their mean. Therefore the central limit theorem can be applied once a large number of samples are obtained, so we can estimate error bounds on 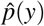 which shrink like 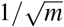. Hence, one can sample until estimates of the evidence are within a specified tolerance. Further, the independence of the sampling procedure allows these computations to be run in parallel and so modern computation hardware can be easily utilised. Of course, having to estimate the likelihood inflates the variance of 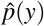, so a larger number of samples are required for 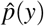 to converge to a given accuracy.

The variance of this estimate is lowest if the sampling density, *q*(*·|y*), is similar to the posterior distribution. Our method requires no parameter inference, but may be made more efficient, especially in higher dimensions, by choosing a sampling distribution similar to the posterior distribution if parameter inference has been performed; for example, the sampling distribution could be a Gaussian with the mean and covariance matrix estimated from MCMC samples [20]. In this paper we do not use MCMC samples to inform the choice of *q*(*·|y*), but this could be done in practice to improve the convergence of estimates.

### 2.2. Sequential importance resampling algorithm for likelihood estimation

Consider a time series, *y* = (*y*_1_*, …, y_T_*), from a partially-observed CTMC. We can express the likelihood at *θ* as

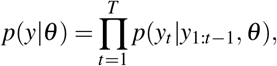

where we adopt the convention that *p*(*y*_1_*|y*_1:0_*, θ*) is taken to mean *p*(*y*_1_*|θ*). Hence, the likelihood can be calculated via a product of the likelihood increments, *p*(*y_t_|y*_1:*t−*1_*, θ*), for *t* = 1*, …, T*. We apply a sequential importance resampling algorithm, which uses importance sampling to estimate the likelihood increments [22, 23]. This algorithm uses resampling at the end of each time increment to lower the variance of likelihood estimates, and remove realisations that were unlikely to have occurred. Throughout this section all probabilities are calculated with respect to some parameter set, *θ* , but this notation is suppressed to make statements concise.

Let *x_t_* denote the state of the partially-observed process at time t. The sequential importance resampling algorithm begins with a set of *n* particles, each particle is associated with an initial state and a weight, {*x*_0_*, w*}^1:*n*^. These initial states are distributed according to *p*(*x*_0_) and the weights begin as 1. Suppose at each iteration of the algorithm we have states 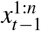 distributed according to particle filter density 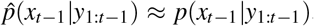 we evolve these over a day according to the importance sampling density, *q*(*·|y_t_, x_t−_*_1_), to obtain realisations 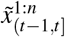 with weights given by

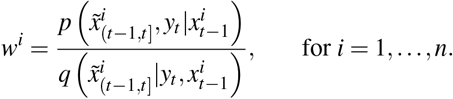

These are computed such that the expected value of the weights are

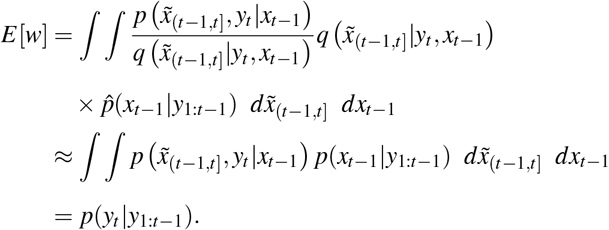

Hence, the mean of these weights provides an approximation of the likelihood increment, *p*(*y_t_|y*_1:*t−*1_); the product of these increments gives an unbiased estimate of the likelihood [24]. Further, resampling states from 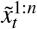 according to normalised weights, 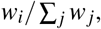 gives a sample, 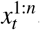, from 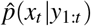. Hence, from the initial conditions we can iterate forwards and recursively estimate the likelihood. Performance of this algorithm can be improved by updating weights over multiple steps and only resampling once the effective sample size drops below a specified threshold [25].

Thus far we have described the general approach of using importance sampling and sequential importance resampling to estimate posterior model probabilities, however, an important aspect of the sequential importance sampling process is the evolution of particles over each day. The difficulty in evolving particles in the cases considered here is that precise observations are made from models with many latent variables, which can make data-augmentation or rejection-sampling approaches slow. In Sections 3.2 and 4.3 we describe the method from [8], which builds upon [26], for generating realisations 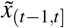 from initial state *x_t−_*_1_ for each of the case studies. The main benefit of this approach is that we use an efficient sampling distribution *q*(*·|y_t_, x_t−_*_1_) which is tractable and generates 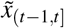 which match observations almost surely, and, are similar to the partially-observed process. This method makes particles match observations by first generating observation times, then generating events between observations and occasionally forcing events to occur or blocking events to ensure feasibility of the process. For example, if we are yet to observe an infection but a recovery would lead to the epidemic ending, the recovery rate is set to 0. Importance sampling in this way allows for estimation of likelihoods associated with rare events – for example it can be easily used to estimate the tails of the likelihood, or just used to sample when observations are unlikely – whereas rejection sampling methods tend to perform poorly as simulations are unlikely to match the data.

The overall approach to model selection here is to: (i) sample from the parameter space using some importance sampling distribution; (ii) for each of those samples estimate the likelihood by sequential importance sampling; (iii) plug this likelihood estimate into Equation (1) to obtain importance weights; and, (iv) use the mean of these weights as an unbiased estimate of the model evidence. We can keep sampling and obtaining estimates of the model evidence until they satisfy some stopping criteria; for example, once credible intervals are of a specified width. Once the model evidence is computed for all candidate models we can multiply these by the prior distribution over the models and normalise to obtain the posterior model probabilities.

## 3. Case study I: Inferring the infectious period distribution

Our first case study uses the importance sampling Bayesian model selection method to infer the appropriate infectious period distribution for an SI(k)R model [27, 28, 29], where symptoms are assumed to coincide with a transition into the infectious class. This study is motivated by influenza outbreaks in which only symptom onset (which is correlated with infection) is observed, and recoveries are not. Here we consider the infectious period to be either exponential, Erlang-2 or Erlang-5 distributed; these represent high, medium and low variance infectious periods respectively. This study aims to answer whether case data at a daily resolution is sufficient for discriminating between these models, how well parameters need to be known in order to discriminate between models, and how much data is required to effectively discriminate between models.

It is known that with final size data the SI(k)R model has a tractable likelihood function [30, 29], which allows for efficient Bayesian model selection. But, it is currently an open question as to whether full temporal data is more effective for model selection as, although there is more information in the data, the parameter space is larger (effectively going from 2 to 3 dimensions to also infer *γ*). Hence, we compare model selection results from the full temporal data with results from final outbreak size data.

The temporal data considered in this paper consists of daily symptom onset counts from completed outbreaks in small populations, which we refer to as households. Final size data is derived from the temporal data by summing over the total number of cases in each household. We let all households be of size 4 for simplicity, though this can easily be extended to allow for a distribution of household sizes. Each outbreak is modelled as a compartmental CTMC, however we only observe a small portion of the epidemic process, so this is a partially-observed CTMC. We assume that all events of symptom onset are observed until the epidemic fades out in the household and set the time of the first observation within each household as 0.

We describe the epidemic model for households in Section 3.1, the importance sampling method for estimating likelihood increments in Section 3.2, the parameters used for the simulation study in Section 3.3, and show results in Section 3.4.

### 3.1. SI(k)R model

For a population of size *N*, the SI(k)R model is a compartmental model that allows each individual to be in one of *k* + 2 compartments: they are either susceptible; infectious in phase *j*, for *j* = 1*, …, k*; or, recovered. Here the different phases of infectiousness have no physical interpretation; they are introduced in order to allow the overall infectious period to be Erlang-k distributed [27, 28, 29]. Let *S*, *I_j_* and *R* denote the number of susceptible, infectious phase *j* and recovered individuals respectively. As these numbers must always be non-negative and sum to *N*, we have the state space

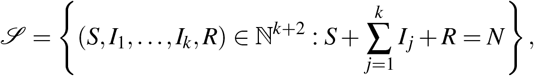

where we take ℕ to contain 0. There are three kinds of transitions for this model: infection; phase change; and, recovery. Infectious individuals of all phases make effective contact with other individuals in the population at rate *β*, and if this contact is with a susceptible individual then that individual becomes infectious, corresponding to a transition into the infectious phase 1 compartment. An infectious phase *j* individual for *j* = 1*, … , k −* 1, moves into the next phase at rate *kγ* ; this rate is chosen such that the infectious period is Erlang-k distributed with mean 1*/γ*. Similarly, an infectious phase *k* individual recovers at rate *kγ* and is no longer able to spread the disease. The transitions and rates associated with changes in the number of individuals in each compartment are given in Table 1.

**Table 1:** Transitions and rates for the SI(k)R model. Only compartments that change are shown, all other compartments remain the same.

| Transition Type | State Change | Rate |
| --- | --- | --- |
| Infection | $(S, I_1) \rightarrow (S - 1, I_1 + 1)$ | $\frac{\beta S \sum_{j=1}^k I_j}{N-1}$ |
| Phase change type j | $(I_j, I_{j+1}) \rightarrow (I_j - 1, I_{j+1} + 1)$ | $k\gamma I_j$ |
| Recovery | $(I_k, R) \rightarrow (I_k - 1, R + 1)$ | $k\gamma I_k$ |

For this study we assume that symptom onset corresponds to the infection transition, and we assume that we observe the number of these transitions at a daily resolution. The model is initialised at the time of the first observations, hence the initial state is (*S, I*_1_*, I*_2_*, …, R*) = (*N −* 1, 1, 0*, … ,* 0).

### 3.2. Importance sampling for SI(k)R outbreaks

Suppose we have observations from *M* outbreaks, *y*^1:*M*^. As these outbreaks occur independently we have a likelihood function which is of the form

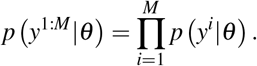

Hence, the likelihood can be estimated via multiplying estimates of likelihoods for each outbreak, *p*(*y^i^|θ*). The sequential importance resampling scheme allows us to calculate each of these iteratively, so we are left to describe how to generate realisations over the day from an initial state, *x_t−_*_1_, in a way that ensures consistency with the observations, and, how to evaluate the importance sampling weights. We do this as per the method of [8].

Suppose for each outbreak we have a dataset *y* = (*y*_1_*, …, y_T_*) where *y_t_* gives the cumulative number of infection events over (*t −* 1*, t*] for *t* = 1*, … , T*; note that this does not include the initial infectious individual in the population. To estimate *p*(*y_t_|y*_1:*t−*1_), we begin by uniformly generating *y_t_* observation times over (*t −* 1*, t*]. These times are ordered, so the joint density is that of *y_t_* order statistics of Uniform(*t −* 1*, t*) random variables, so we initialise the importance weights by *w* = 1*/y_t_* !. Then, beginning from time *τ* = *t −* 1 in state,

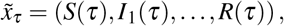

determined by the final state of the previous iteration, we generate events between observation times. However, we only allow phase change or recovery transitions to occur, as the observations (and hence infections) have already been generated. Let *a* be a vector of rates,

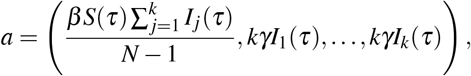

as per Table 1, associated with state 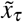 We consider a process with modified rates *b*, given by

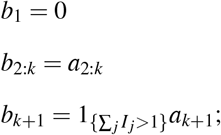

where 1_{*A*}_ is an indicator function which takes the value 1 if logical statement *A* holds or 0 otherwise. The modified rates are constructed so that no further observations (infections) can occur, as *b*_1_ = 0, and no recoveries can occur if it would lead to epidemic fade-out. Let *τ^ʹ^* denote either the next observation (or if there are no further observations over the day let *τ^ʹ^* = *t*). We sample an Exponential(∑ *_j_ b j*) candidate time increment, Δ*τ*. Then one of three kinds of updates occur:

(i) if *τ* + Δ*τ < τ^ʹ^* we generate an event at time *τ* + Δ*τ* and let the event be of type *i* with probability *bi/* ∑ *_j_ b _j_*, the importance weight is updated to

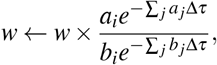 we update the time *τ ← τ* + Δ*τ*, and we update the state to 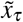 according to transition type *i*;
(ii) if *τ* + Δ*τ > τ^ʹ^* and *τ^ʹ^* ≠ *t* no event occurs in (*τ, τ^ʹ^*) and the next event is an observation, so we update the weights to

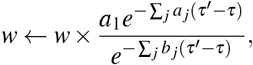 we update the time, *τ ← τ^ʹ^*, and update the state 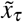 according to an infection transition; or, if
(iii) *τ* + Δ*τ > τ^ʹ^* and *τ^ʹ^* = *t* then no event occurs before the end of the day and the weights are updated to

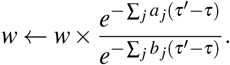

We repeatedly recalculate the rates and make updates until an update of type (iii) occurs, at which time the process ends for this time increment.

Once all particles have gone through this process the weights are averaged to form an approximation of *p*(*y_t_|y*_1:*t−*1_) and the states are resampled, using systematic resampling [31], according to the normalised weights. These samples provide initial states for calculating the next likelihood increment. Once the last observation in the time series has been generated, we generate from a modified process where *b* = (0*, a*_2:*k*+1_) and update weights according to (i) until the epidemic dies out; this allows us to calculate the portion of the likelihood associated with the assumption that the epidemic died out after our last observation.

### 3.3. Implementation

We simulate 50 temporal and final size data sets of multiple completed outbreaks in households of size 4 with parameters (*β, γ*) = (0.933, 2*/*3) under each of the three models. We use 500 particles per likelihood calculation, as this is found to be sufficient for low variance likelihood estimates. Our implementation uses the prior distribution as the importance sampling distribution over the parameter space, *q*(*θ|y*), however a more efficient sampler could be chosen if PM-MCMC is performed before model selection [20]. We begin by sampling 500 points from *q*(*θ|y*) and continue to sample in batches of 500 samples from the parameter space until 95% credible intervals of the model evidence are non-overlapping. The initial 500 samples are such that the central limit theorem can provides estimates of precision of model evidence with small bias. Sampling in batches allows for sample weights to be calculated efficiently in parallel before the precision of the evidence is calculated again. The stopping criterion is chosen so that we can accurately choose the most appropriate model; in practice other stopping criteria could be used to calculate model evidence to a given precision. Implementing a stopping criterion in this way ensures that the number of particles for point estimates of the likelihood only effects the run time of the algorithm, not the precision of model evidence estimates. We calculate posterior model probabilities using data from 50, 100 and 150 complete household outbreaks. For each of these data sets we consider two cases: where the mean infectious period and the reproduction number are known to a high level of accuracy *a priori*; and, where the prior distribution on model parameters is relatively uninformative. We refer to these as the assumption of tight priors and loose priors, respectively. We set the mean of the tight priors to their true value; we suppose that 1*/γ* has a gamma distributed prior with mean 3*/*2 and variance 0.01, and that *β/γ* has a Uniform(0.933 *×* 3*/*2 *−* 0.03, 0.933 *×* 3*/*2 + 0.03) prior. For the loose priors we suppose that 1*/γ* has a gamma distributed prior with mean 2 and variance 0.75, and we assume that *β/γ* has a Uniform(1, 2) prior; which is a typical range of *β/γ* for influenza. Note the likelihood estimates can have high variance in regions of the parameter space where the rate associated with observed events is large [8]. This issue is avoided as the prior distributions ensure that parameters only have support in places away from these values. In the simulation study the relatively low variance of likelihood estimates allowed the sample variance of the weights to remain low enough for convergence to occur.

### 3.4. Results

Our results are given in terms of box plots of the difference in posterior model probabilities of the true model and other candidate models in Figures 1 and 2, and in terms of the proportion of times the correct model was identified (having the highest posterior model probability) in Table 2. If the box plots are near one it means the posterior model probability for the correct model is nearly one; values that are negative represent times at which the other candidate models have higher posterior model probabilities.

**Figure 1:**
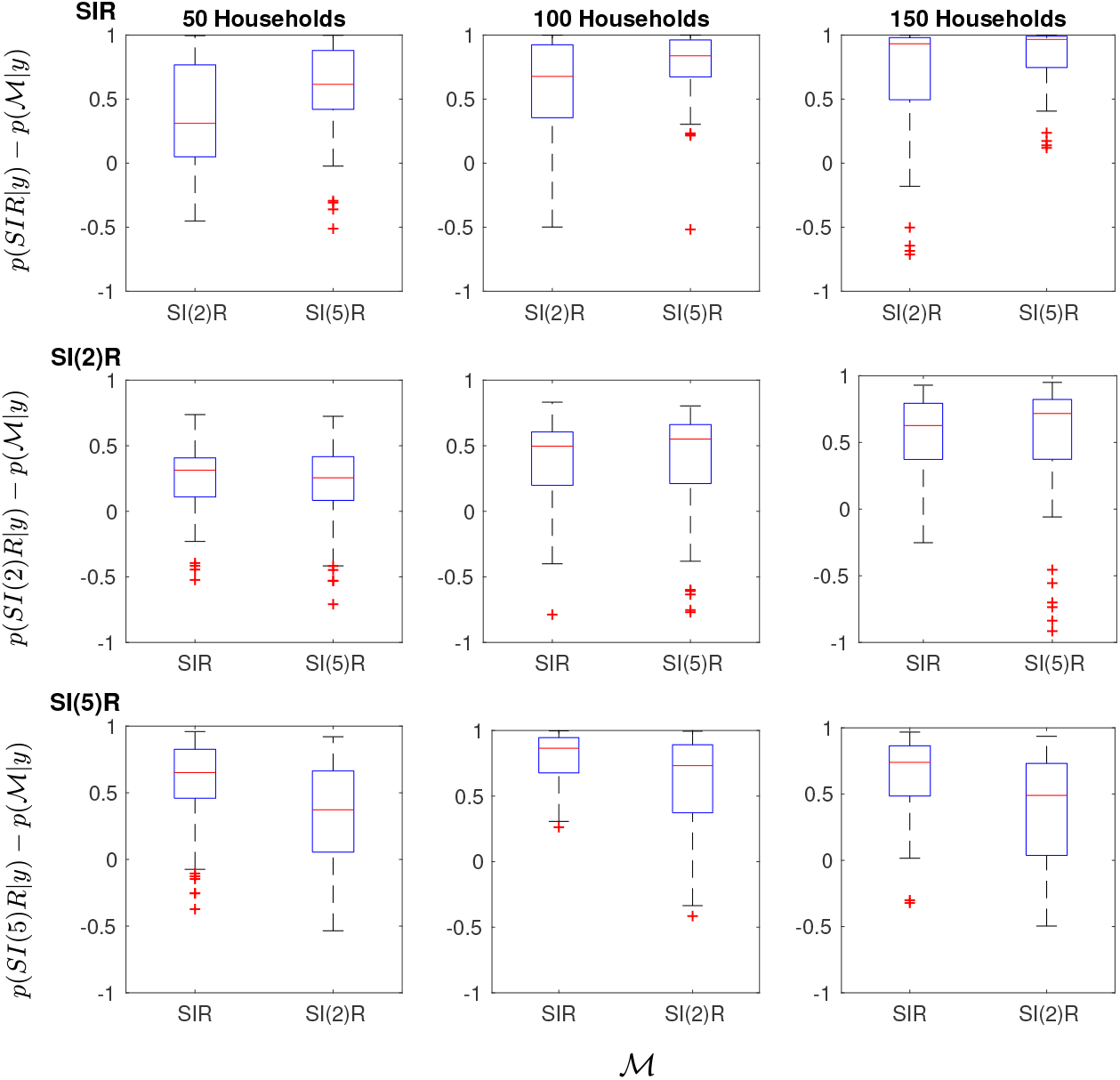
Box plots of the difference in posterior model probability of the true model and the other candidate models for the SI(k)R models with tight priors based on 50 simulated data sets. For example, in the upper left panel, boxes on the left and right of are made using 50 estimates of *p*(SIR*|y*) *− p*(SI(2)R*|y*) and *p*(SIR*|y*) *− p*(SI(5)R*|y*) respectively. Rows from top to bottom show results from data sets generated from the SIR, SI(2)R and SI(5)R models. Columns from left to right represent data sets containing 50, 100 and 150 independent outbreaks in households.

**Figure 2:**
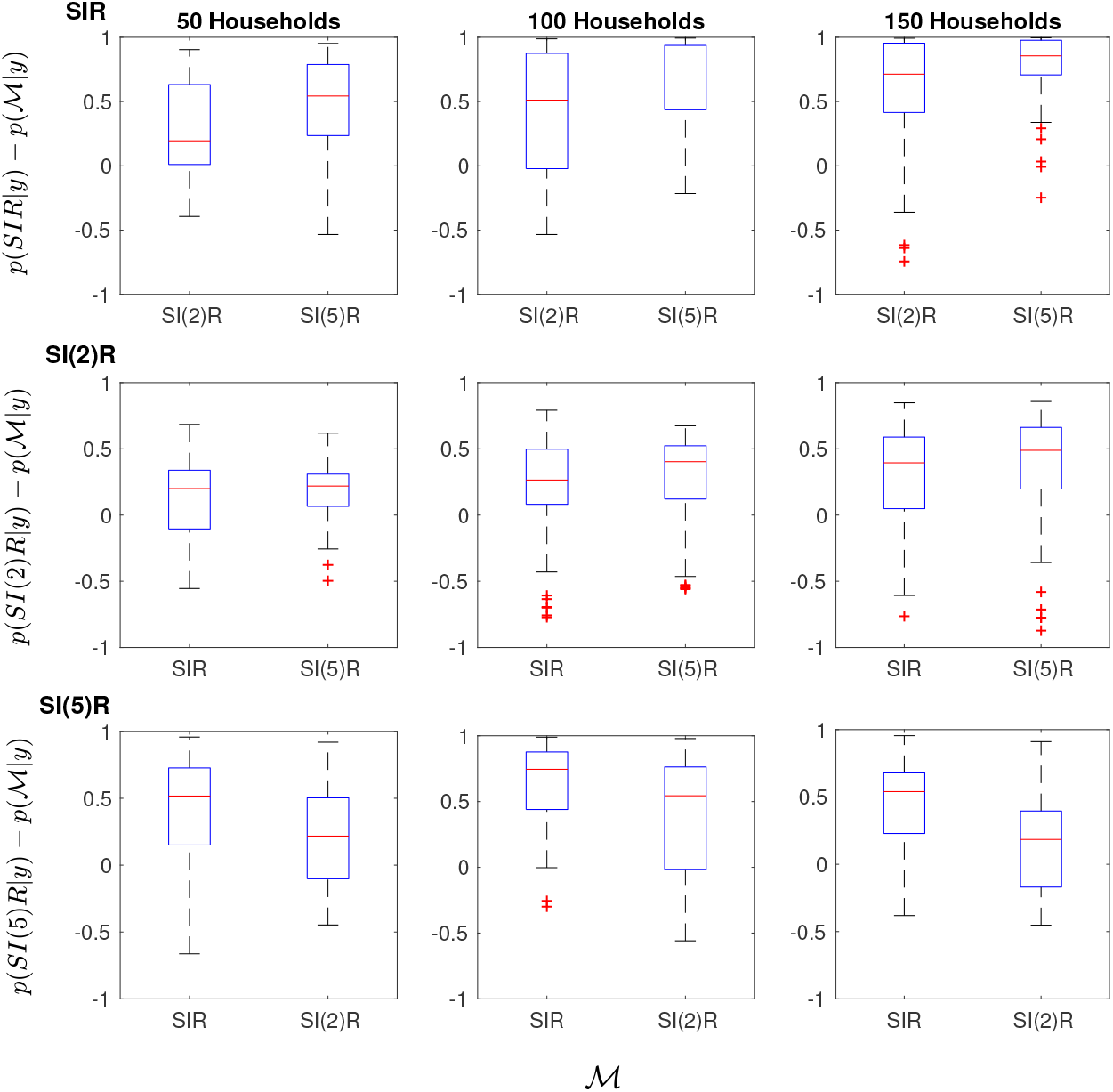
Box plots of the difference in posterior model probability of the true model and the other candidate models for the SI(k)R models with loose priors based on 50 simulated data sets. For example, in the upper left panel, boxes on the left and right of are made using 50 estimates of *p*(SIR*|y*) *− p*(SI(2)R*|y*) and *p*(SIR*|y*) *− p*(SI(5)R*|y*) respectively. Rows from top to bottom show results from data sets generated from the SIR, SI(2)R and SI(5)R models. Columns from left to right represent data sets containing 50, 100 and 150 independent outbreaks in households.

**Table 2:** The proportion of times the model that generated the data corresponded to the highest posterior model probability estimate from 50 data sets. Data sets were generated from each of the SIR, SI(2)R and SI(5)R models with 50, 100, and 150 outbreaks. These data sets were analysed using the full temporal data and the final size data under the assumption of loose or tight priors.

|  |  | Loose priors |  |  | Tight priors |  |  |
| --- | --- | --- | --- | --- | --- | --- | --- |
| Number of outbreaks |  | 50 | 100 | 150 | 50 | 100 | 150 |
| Temporal data | SIR | 0.76 | 0.72 | 0.82 | 0.78 | 0.86 | 0.86 |
|  | SI(2)R | 0.44 | 0.6 | 0.62 | 0.58 | 0.72 | 0.8 |
|  | SI(5)R | 0.68 | 0.74 | 0.86 | 0.8 | 0.84 | 0.96 |
| Final size data | SIR | 0.7 | 0.76 | 0.74 | 0.72 | 0.84 | 0.82 |
|  | SI(2)R | 0.2 | 0.4 | 0.52 | 0.36 | 0.56 | 0.74 |
|  | SI(5)R | 0.6 | 0.58 | 0.64 | 0.62 | 0.74 | 0.78 |

For tight priors, the correct model was most often associated with the highest posterior model probabilities for each of the models considered (Table 2). Note that the lowest box in the panels on Figures 1 and 2 always corresponds to an adjacent model. This fits with the intuition that adjacent models are most often misidentified as each other, that is, models with more similar infectious period distributions are more difficult to discriminate. The SI(2)R model is most often misidentified; this agrees with intuition as this model’s infectious period has a shape parameter between the other two (Table 2). Results were similar under loose priors, (Figure 2), however the box plots tended to have a larger range, showing that there was less certainty in the correct model. There was also very little difference in the proportion of correct times the model was identified under loose and tight priors (Table 2). Even if the parameters are not well known, with multiple completed outbreaks (from small populations), infectious case data can be used to distinguish between these models, but the SI(2)R model is the most difficult to identify. Of the 900 runs, 744 estimates had converged in the first 500 samples, another 115 had converged in less than 10,000 samples and all of the samples converged in under 91,000 samples. Slow convergence occurs when two models are nearly equally likely, in this case it may not be relevant as to whether credible intervals are non-overlapping. This could be avoided by implementing a stopping rule which ensures posterior model probabilities are either non-overlapping or within a specified tolerance. Estimates based on tight priors converged at least as fast as those with loose priors in all but 13 cases.

We compare the results with those obtained by only considering the final size data of the same simulated data sets. Here sampling over the parameter space is the same, however, now the likelihood function is calculated exactly as in [29]. For all cases except for the SIR model with loose priors and 50 outbreaks, one-sided Wilcoxin signedrank tests at the 95% level show that the posterior model probabilities of the true model are statistically significantly higher when the full temporal data is used. The proportion of times the correct model was identified from final size data is given in Table 2. Interestingly, for the SI(2)R model with loose priors and 50 outbreaks the correct model was identified less often than if we had uniformly randomly guessed between the models. This model also saw the biggest improvement from temporal data; the proportion of times the correct model was identified more than doubled. We find that in all cases, except for the SIR model with loose priors and 100 outbreaks, using the full temporal data increased the proportion of times that the correct model was identified. These results show that the full temporal data sets are useful for performing model selection, even though they are more computationally intensive to work with.

As the runtimes of the algorithm were reasonable and we chose a number of particles for likelihood estimates to ensure that estimates of precision of the model evidence had low bias, the number of state particles were not chosen to optimise efficiency. If one had a fixed computational budget and required estimates of model evidence to be as accurate as possible it would be important to make the algorithm as efficient as possible. As such, we have included a comparison of the coefficient of variation of model evidence estimates against the number of particles used in likelihood calculations, *n*, in Figure 3. For three data sets from the SI(2)R model we ran the model selection algorithm for a fixed computation time with differing numbers of particles. The sample weights were used to estimate the coefficient of variation as 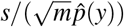, where *m* is the number of samples from the parameter space, *s* is the sample standard deviation of weights, and 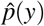 is the mean of the weights. We find that the smallest coefficient of variation is achieved at *n* = 125 for one data set and *n* = 75 for the other two. This indicates that the algorithm would run more efficiently with *n* around 100. As run time per sample is linear with respect to *n* there will be 5 times the number of samples per unit time if *n* were decreased to 100.

**Figure 3:**
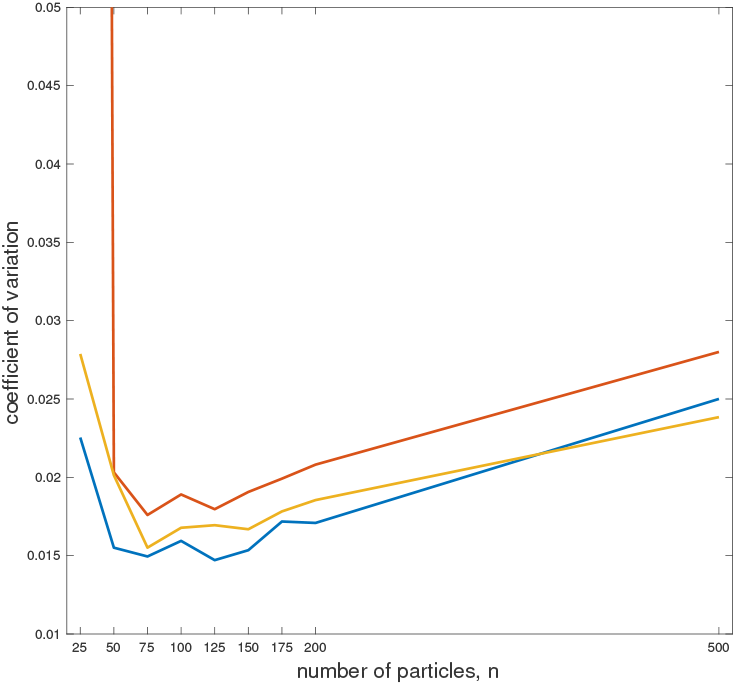
The coefficient of variation of the model evidence estimate against *n*, the number of particles used for likelihood calculations. For three SI(2)R data sets we estimate the coefficient of variation after running the model selection algorithm for 24 hours. The coefficient of variation is estimated as 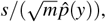, where *m* is the number of samples from the parameter space, *s* is the sample standard deviation of the weights, *Z*_1_, …, *Z_m_*, and 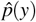 is the mean of the weights.

## 4. Case study II: Inferring symptom onset relative to infectiousness

This study considers inferring the time of infectiousness relative to symptom onset, where infectiousness could either be pre-symptomatic (Pre), coincidental-symptomatic (Co) or post-symptomatic (Post). For example, influenza may have symptoms that largely coincide with infectiousness [32], SARS is largely infectious post-symptom onset [33] and HIV is infectious pre-symptom onset [3]. We consider diseases that have a lag between exposure of individuals and infectiousness, specifically an SE(2)I(2)R model, which is detailed in Section 4.1. This kind of model has been used in previous work on inference using early outbreak data; it has realistic features, such as nonexponential exposed and infectious periods, while being simple enough for inference [34, 35, 36, 37, 7]. We model the observations of symptom onset as either a transition into an exposed, infectious or recovered state for the Post, Co and Pre models respectively (Figure 4).

**Figure 4:**
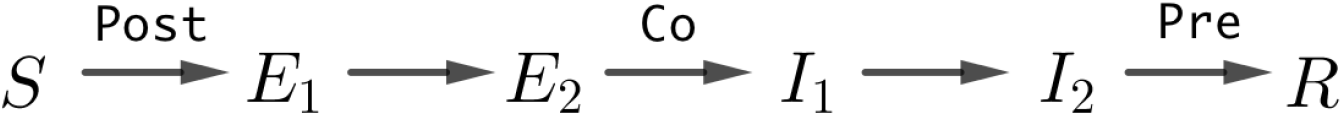
A diagram showing how we relate post-symptomatic (Post), coincidental-symptomatic (Co) and pre-symptomatic (Pre) infections to transitions in the SE(2)I(2)R model.

### 4.1. SE(2)I(2)R model

For a population of size *N*, the SE(2)I(2)R model is a compartmental model that allows each individual to be in one of six compartments: they are either susceptible; exposed phase 1 or 2; infectious phase 1 or 2; or, recovered. The key differences between this model and the SI(k)R model is that the infectious period is assumed to be Erlang-2 distributed and there is an Erlang-2 distributed lag between being exposed to the disease and being able to spread it. Let *S*, *E_i_*, *I_j_* and *R* denote the number of susceptible, exposed phase *i*, infectious phase *j* and recovered individuals respectively. Again, as these numbers must always be non-negative and sum to *N*, we have state space

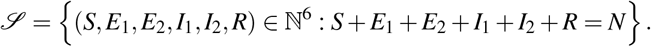

There are five kinds of transitions: exposure; exposed phase change; infectious; infectious phase change; and, recovery. These transitions are similar to those described for the SI(k)R model and are shown in Table 3.

**Table 3:** Transitions and rates for the SE(2)I(2)R model. Only compartments that change are shown, all other remain the same.

| Transition Type | State Change | Rate |
| --- | --- | --- |
| (1) Exposure | $(S, E_1) \rightarrow (S - 1, E_1 + 1)$ | $\frac{\beta S(I_1 + I_2)}{N - 1}$ |
| (2) Exposed phase change | $(E_1, E_2) \rightarrow (E_1 - 1, E_2 + 1)$ | $2\sigma E_1$ |
| (3) Infectious | $(E_2, I_1) \rightarrow (E_2 - 1, I_1 + 1)$ | $2\sigma E_2$ |
| (4) Infectious phase change | $(I_1, I_2) \rightarrow (I_1 - 1, I_2 + 1)$ | $2\gamma I_1$ |
| (5) Recovery | $(I_2, R) \rightarrow (I_2 - 1, R + 1)$ | $2\gamma I_2$ |

We assume that symptom onset corresponds to a transition into either the exposed phase 1, infectious phase 1 or recovered class for the Post, Co and Pre models respectively, as shown in Figure 4. Hence, the process has an initial state that is either

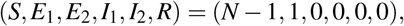

for exposure observations,

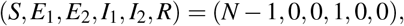

for infectious observations, or a stochastic initial state for the recovery observations; this last case is discussed in the following section.

### 4.2. Importance sampling for initial state generation

For the SE(2)I(2)R model, if we observe the daily number of recovery transitions, then a population with a single exposed individual initially may be in one of several states at the time of the first observation, as multiple exposures may have occurred before the first recovery. If we see no secondary transmission in the household we can calculate the likelihood exactly as

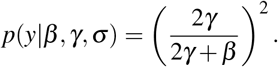

If the epidemic does not die out after the initial infection we can efficiently generate weighted initial states of the process via importance sampling.

If our data set for the household has daily observation vector 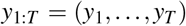 we know that each type of transition can occur at most 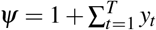 times and that recovery cannot occur if it would lead to epidemic fade out; note that the plus 1 is because the initial condition is not included in *y*_1:*T*_. So if the SE(2)I(2)R process has rate vector

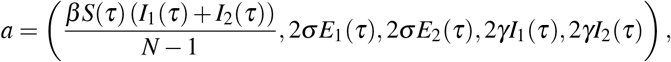

as per Table 3, we consider a modified process with rates that never lead to inconsistencies in our data. The modified rates are

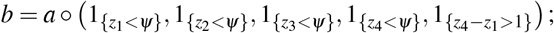

where ‘*◦*’ denotes an elementwise product, *z_i_* denotes the cumulative number of individuals that entered the *i*th compartment, and 1*_{·}_* denotes the indicator function. The modified process is used to generate transitions until an observation occurs; the bias from simulating from this modified process is corrected for by using importance sampling weights.

The process begins with particles in state

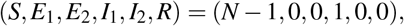

with importance weight *w* = 1. Each particle makes transitions according to probabilities *b/* ∑ *b*, and with each transition the particles weight is updated to

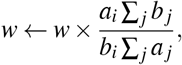

where *i* is the type of the transition that occurred (type is numbered as per Table 3). When the first recovery (transition type 5) occurs the process ends. Once we have a set of particles distributed according to the initial state they are resampled according to the normalised importance weights to give initial particles for the sequential importance resampling algorithm.

### 4.3. Importance sampling for SE(2)I(2)R outbreaks

The importance sampling scheme for the SE(2)I(2)R model is similar to the SI(k)R model except that the modified rates are slightly different, and, at times, we need to force certain events to occur to ensure feasibility of samples [8]. Again, we uniformly generate observation times over (*t −* 1*, t*] and from time *t −* 1 generate rates between observations according to a modified process with rate vector, *b*. We set modified rates *b _j_* = *a _j_* for all *j* that correspond to rates which are not set to 0 (those set to 0 will be specified in this section). For the Post, Co and Pre models we set the modified rates *b*_1_ = 0, *b*_3_ = 0 and *b*_5_ = 0 respectively. We also set *b*_5_ to 0 if a recovery would lead to epidemic fadeout without the correct number of observations occurring. For these models we may also need to force events to occur to ensure that the particle generated, 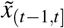 is a feasible realisation from an SE(2)I(2)R process. For example, for the Co model, if there are no exposed phase 2 individuals in the population and an observation is yet to occur, an exposed phase change would need to occur. If there were also no exposed phase 1 individuals then an exposure time would need to be generated before the exposed phase change; we refer to these kinds of events, and observation events, as *forced events*. More generally, if the particle is a realisation generated up to time *τ*, 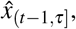, where the next forced event occurs at time *τ^ʹ^* and would lead to an infeasible realisation, we generate a new forced event which would allow the next forced event to be feasible. The next forced event is chosen by proposing transitions further back in the chain, or forcing an infectious event (transition type 3), until an event with a positive rate is proposed. The order in which to propose events is shown via a flowchart in Figure 5. For other CTMC’s this can be done by proposing forced events according to a logic tree which is model specific. If we need to generate a forced event of type *i*, we generate an inter-arrival time, *s*, according to a TruncExp(*a_i_, τ^ʹ^ − τ*) distribution, set *b_i_* = 0, set the next forced event as type *i* at time *τ^ʹ^* = *τ* + *s*, where TruncExp(*a, t*) refers to the truncated exponential distribution with rate *a*, truncated to [0, t]. The truncated exponential distribution is chosen as it has appropriate support and generates events with a similar distribution to the true process. The weights are updated according to

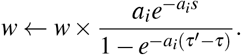

**Figure 5:**
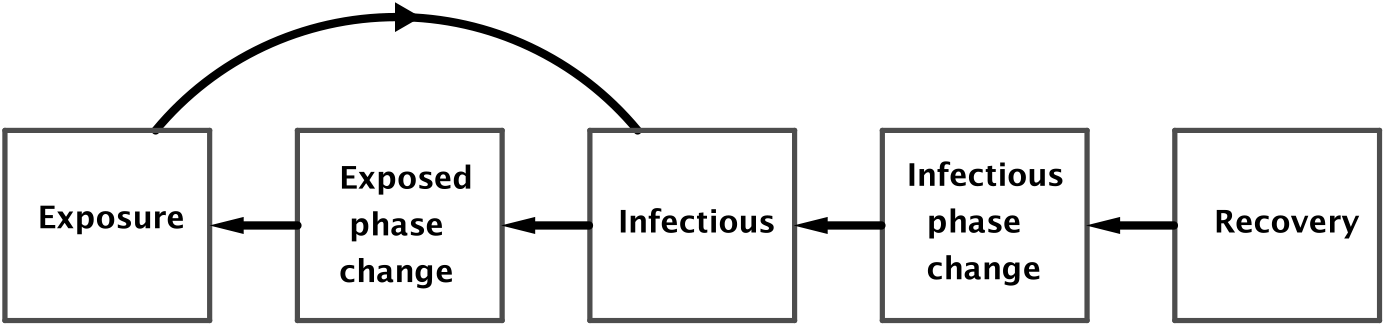
A flow chart showing the order in which transitions are proposed as forced events in order to ensure feasibility of the particle. Starting from the type of the next forced event, propose a new forced events according to arrows in the flow chart.

If no more forced events are necessary we continue to propose candidate events from time *τ* according to the new modified rates; as in Section 3.1, we let these occur, move to the next forced event, or move to the end of the time increment as per (i), (ii) and (iii) respectively. Once the cumulative number of events of type *i* equals the number of observations in total we set *b_i_* = 0; as the epidemics are completed within households the final state must be (*N − ψ,* 0, 0, 0, 0*, ψ*), so no event can occur more than *ψ* times.

For more details and a more general description of the importance sampling scheme see [8].

### 4.4. Implementation

We simulate 20 data sets of multiple completed outbreaks in households of size 4 with parameters (*β, σ, γ*) = (0.933, 0.5, 2*/*3) under each of the three models and calculate posterior model probabilities with data from 50, 100 and 150 completed household outbreaks. In this case we used 2000 particles per likelihood calculation; a larger number of particles were chosen because the sampling distribution is less like the true process, due to the possibility of needing to force events. Our implementation uses the prior distribution as the importance sampling distribution over the parameter space, *q*(*θ|y*). We first sample 1000 points from the parameter space and continue to sample from the parameter space until 95% credible intervals of the model evidence are non-overlapping. We consider inference based upon tight and loose priors on all model parameters. We set the mean of the tight priors to their true value; we suppose that 1*/γ* has a gamma distributed prior with mean 3*/*2 and variance 0.01, similarly we assume that 1*/σ* has a gamma distributed prior with mean 2 and variance 0.01 and we assume that *β/γ* has a Uniform(0.933 *×* 3*/*2 *−* 0.03, 0.933 *×* 3*/*2 + 0.03) prior. For the loose priors we suppose that 1*/γ* and 1*/σ* have gamma distributed priors with mean 2 and variance 0.75 and we assume that *β/γ −* 1 has a gamma distributed prior with mean 1 and variance 0.75.

### 4.5. Results

Our results are given in terms of box plots of the difference in posterior model probabilities of the true model and other candidate models in Figures 6 and 7, and in terms of the proportion of times the correct model was identified in Table 4. The Pre model is the most difficult model to identify with data on 50 outbreaks. With data on 150 outbreaks the correct model was selected every time except for data generated from the Co model. The boxes in Figure 6 are all situated near 1, indicating that with tight priors, when only 50 households are infected, we are usually certain of which model is the true model. By the time 150 households are infected effectively all posterior model probabilities are close to 1, so we are almost always certain of which model is the true model. The boxes in Figure 7 are situated lower than those in Figure 6, indicating that loose priors reduces the certainty of the correct model. However, each model is easily identifiable whether or not the priors are informative and by the time 150 households have had completed outbreaks the correct model is identified in almost all cases. Figure 7 also shows that even with loose priors the posterior model probabilities for the correct model are almost always near 1 for Post and Pre models once 150 households are infected. It also shows that the Co model tended to be chosen with less certainty when priors were loose, with one outlier choosing an incorrect model with posterior model probability near 1. Of the 360 runs, 336 converged in the first 1000 iterations and all 360 runs converged within 5000 iterations; the fast convergence is likely due to easy identifiability of the three models.

**Figure 6:**
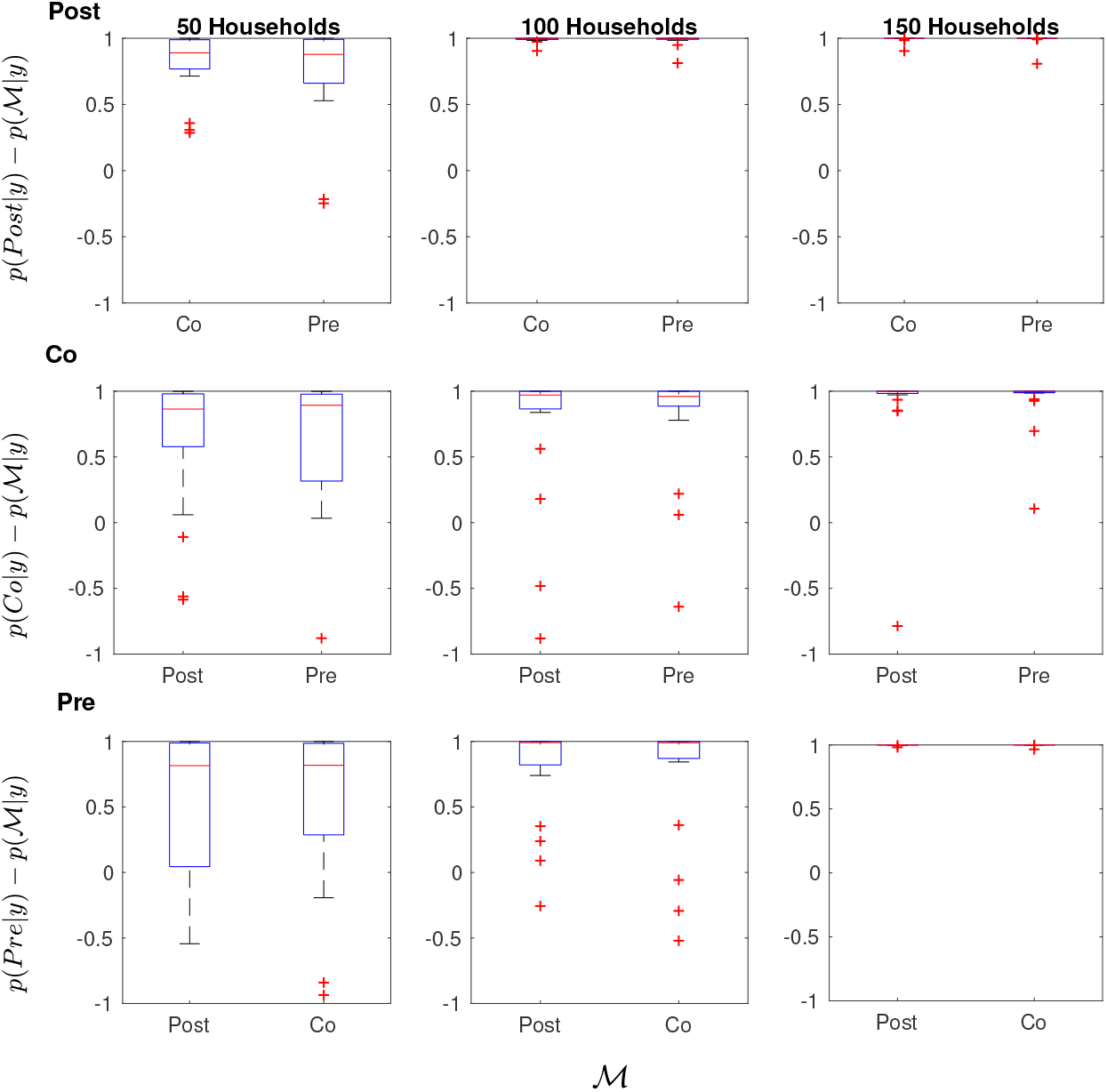
Box plots of the difference in posterior model probability of the true model and the other candidate models for the SE(2)I(2)R observation models with tight priors based on 20 simulated data sets. For example, in the upper left panel, boxes on the left and right of are made using 20 estimates of *p*(Post*|y*) *− p*(Co*|y*) and *p*(Post*|y*) *− p*(Pre*|y*) respectively. Rows from top to bottom show results from data sets generated from the Post, Co and Pre models. Columns from left to right represent data sets containing 50, 100 and 150 independent outbreaks in households.

**Figure 7:**
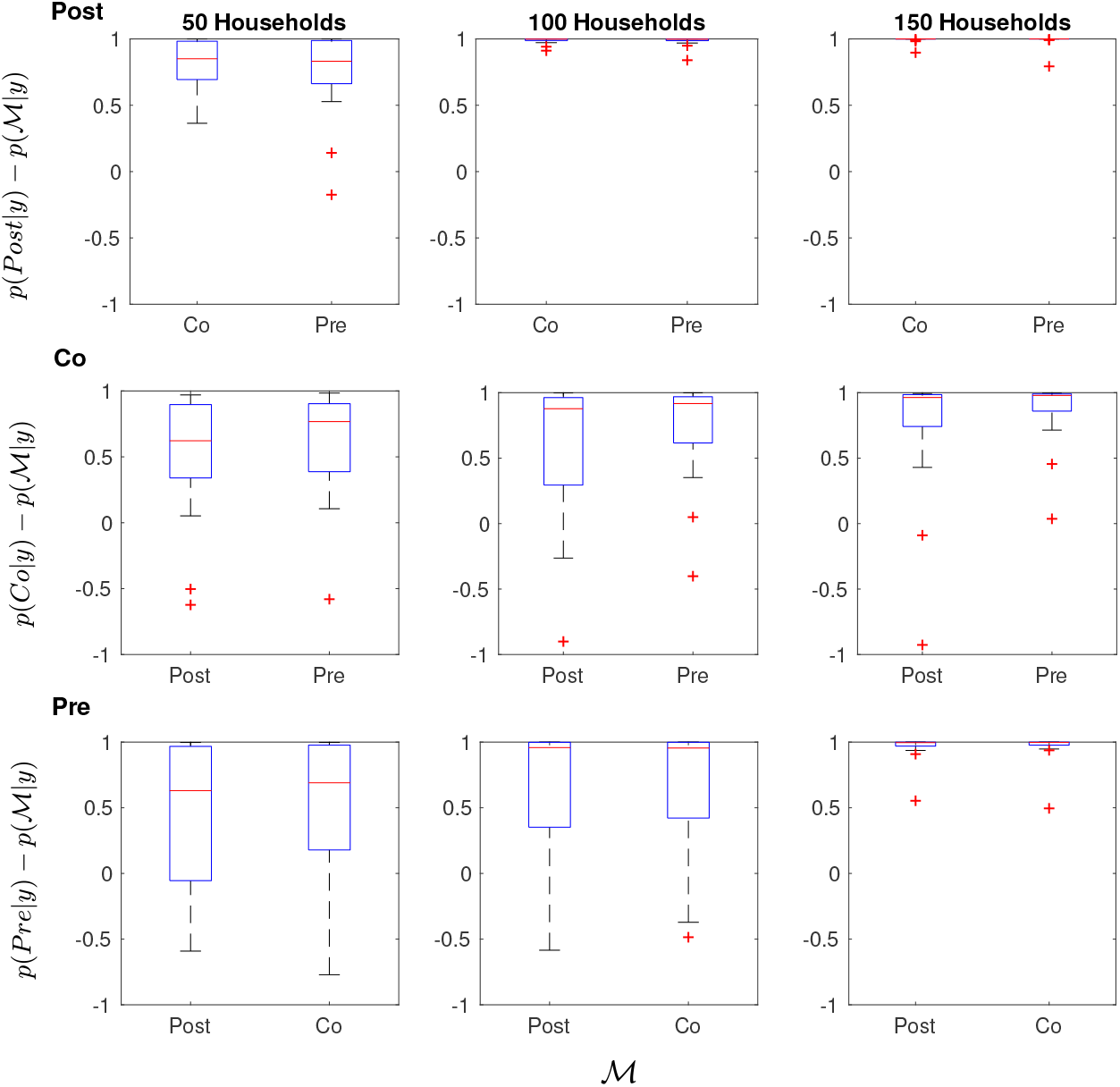
Box plots of the difference in posterior model probability of the true model and the other candidate models for the SE(2)I(2)R observation models with loose priors based on 20 simulated data sets. For example, in the upper left panel, boxes on the left and right of are made using 20 estimates of *p*(Post*|y*) *− p*(Co*|y*) and *p*(Post*|y*) *− p*(Pre*|y*) respectively. Rows from top to bottom show results from data sets generated from the Post, Co and Pre models. Columns from left to right represent data sets containing 50, 100 and 150 independent outbreaks in households.

**Table 4:** The proportion of times the model that generated the data corresponded to the highest posterior model probability estimate from 50 data sets. Data sets were generated from each of the Post, Co and Pre models with 50, 100, and 150 outbreaks. These data sets were analysed under the assumption of loose or tight priors.

| Number of outbreaks | Loose Priors |  |  | Tight Priors |  |  |
| --- | --- | --- | --- | --- | --- | --- |
|  | 50 | 100 | 150 | 50 | 100 | 150 |
| Post | 0.95 | 1 | 1 | 0.9 | 1 | 1 |
| Co | 0.85 | 0.8 | 0.9 | 0.8 | 0.85 | 0.95 |
| Pre | 0.7 | 0.8 | 1 | 0.75 | 0.8 | 1 |

## 5. Discussion

This paper has introduced an exact method for Bayesian model selection which used importance sampling for estimating the likelihood function as well as for estimating the evidence. The novelty of this method is the use of an efficient importance sampling scheme ideal for partially-observed state space models used to estimate the likelihood function [8]. Implementation of this scheme has some overhead compared to Doob-Gillespie simulations; however, by construction all simulations agree with the observed data, hence all strictly contribute to the likelihood estimate. This is in sharp contrast to rejection-sampling approaches which do not force samples to fit with observations, such as in approximate Bayesian computation [38] or the Alive particle filter [39, 19]. Hence, it is able to greatly improve computational efficiency, particularly in estimating the tails of the likelihood. This is also an *exact* method, in that it computes unbiased estimates which converge to the target as the number of iterations increases. In contrast, other methods may not be guaranteed to converge; for example, approximate Bayesian computation, which accepts samples that do not perfectly agree with data. Further, methods may have an exact implementation, but in practice approximations are needed for computational feasibility; for example, the Alive particle filter [39, 19] can be implemented as an exact method but may require too many simulations to be practical, so the likelihood function is set to 0 in the tails instead.

If it is possible to evaluate conditional densities, a different approach is to directly calculate the evidence after use of a Gibbs sampler [40, 41]. This is problematic in that typically the Gibbs sampler for partially-observed CTMCs will require data augmentation [42, 6], making mixing slow and convergence problematic when the amount of missing data to be imputed becomes large [26, 43, 6]. Further, the conditional densities are usually only analytically tractable if priors are conjugate. In [20] data-augmentation is used to estimate the evidence, but this required a time discretisation of the model so that the forward-filtering backwards-sampling algorithm could be applied to sample from the latent process. If the model is not discretised and it is not possible to sample exactly from the latent process, data-augmentation may still be applied. However, for each point estimate of the likelihood function this requires imputing latent variables until the process reaches stationarity. Imputation is undesirable as convergence can be slow for models with high dimensional latent processes and likelihood estimates may still have a high variance. Our method circumvents this difficult imputation by instead marginalising over the latent variables in estimating the likelihood. A weakness of our approach is that we are currently restricted to Markovian models where a single event type is observed whereas data-augmented approaches are much more flexible. Current research is looking at how our method can be extended to situations where multiple types, and combinations, of events are observed.

Other methods for Bayesian model selection include reversible-jump MCMC (RJ-MCMC) [5], nested sampling [44, 45] and sequential Monte Carlo-squared (SMC^2^) [46, 19, 47]. RJ-MCMC is an algorithm in which the model is considered as one of the parameters to be sampled [5]. One issue with such an approach is that if models are non-nested it can be difficult to make sensible proposals between the different parameter spaces, particularly if data-augmentation techniques are used to address unobserved transitions [20]. Hence, at times, proposals are accepted with low probability and the mixing of the algorithm can be prohibitively slow. Nested sampling is an approach to model selection which does not require MCMC samples, however it requires a tractable likelihood [44, 45], which is uncommon for most dynamical mechanistic models. SMC^2^ is an alternative that jointly infers model parameters and the posterior model probabilities by using particle filters in the parameter space and in the state space [46, 19, 47]. As noted by [19], the estimates that allow for model selection may have large Monte Carlo error, but use of importance sampling post-SMC^2^ can be an effective way of reducing this error. During the revision process we became aware, by a personal communication, of a similar method for Bayesian model selection for partially-observed continuous-time Markov chains using importance sampling [48]. The paper uses an alive particle filter to estimate the likelihood function. The alive particle filter, in practice, introduces bias into likelihood estimates in the tails [19, 48]. The particle filter used in this paper does not introduce any bias into the likelihood function. Further, we consider different examples from epidemiology, which are important for understanding emerging infectious diseases.

Other than in [48] and this paper, the method of importance sampling for estimating the evidence has typically only been considered for cases where the likelihood function is known [4, 21], for inference on continuous-state models [21], where data augmentation is used to estimate the likelihood function [20, 21], or in conjunction with another method which is not suitable for processes with highly variable observations [19]. This sort of implementation is inefficient for models with high dimensional parameter spaces. One way to overcome this is by using particle-marginal MCMC steps to inform a sampling distribution over the parameter space prior to model selection [20].

In this paper we described the general method used but made no attempt to optimise the algorithm. We found that, in cases where posterior model probabilities were close, it may take many samples before estimates have non-overlapping credible intervals. Performance could be improved by implementing a stopping criterion where sampling occurs until either evidence estimates are non-overlapping or are within a specified tolerance. For the simulation study we chose a large number of state particles, *n*, for likelihood calculations to avoid needing a large number of samples from the parameter space. The choice of the number of particles for estimating the likelihood was chosen for reasonable performance, rather than optimal performance. There is potential to improve efficiency by informing the number of particles based on optimality criteria, such as minimising the coefficient of variation of evidence estimates after a fixed amount of computation. We estimated the coefficient of variation after running the model selection algorithm for a fixed computation time on three SI(2)R data sets with differing numbers of particles. We found that the coefficient of variation was minimised at *n* = 125 in one case and the other two were minimised at *n* = 75. If we had chosen *n* = 100, say, the runtime of the algorithm would be reduced. As run time per weight calculation is linear in *n*, this would represent a five-fold increase in the number of samples per unit time, though the variance of weight estimates would be larger. We also note that the models considered in this paper are likely to give rise to household outbreak data sets that are equivalent, so computational gains could be made by reusing likelihood estimates for these data sets, as in [7]. However, these steps were not considered in this paper as the computational benefit is model specific and is decreased when considering a slightly more complicated example, such as a model with heterogeneous household sizes. Whereas, the current implementation is versatile enough that it will readily run on models with heterogeneous household sizes.

A number of other approaches to model selection exist. Competing models are commonly discriminated using information criteria which are based on maximum likelihood estimates, such as AIC [49], AICc [50], BIC [51] or DIC [52]; note none of these account for prior information about model parameters, they depend on asymptotic results or depend on distributions being approximately Gaussian. Further, interpretation of these quantities is non-intuitive other than that they represent a function of maximum likelihood values which are penalised for each parameter in the model, and comparison of Bayesian model selection and DIC found DIC to be unreliable [53].

Our method was applied to two case studies: inferring the shape of the infectious period distribution of an SI(k)R, model and inferring the time of symptom onset relative to infectiousness for an SE(2)I(2)R model. In each of the studies we considered daily symptom onset data from completed outbreaks in multiple households. The first study showed that the data was sufficiently informative to select the correct model much more often than by randomly guessing. The study also showed that temporal data was better able to choose an appropriate model than the final size data alone, although at a higher computational cost. For example, from data generated from 150 outbreaks from the SI(5)R model with loose priors the proportion of times the correct model was identified was 0.64 from final size data compared to 0.86 from the full temporal data. Although using the full temporal data is more computationally expensive, when run on multiple CPUs the runtime is divided by the number of CPUs used, indicating that the full data sets should be used if the computational facilities are available. The SI(2)R model was the most difficult model to select correctly, intuitively this seems to be because the disease has an infectious period distribution that has a variance between the other two models. It is worth noting that the posterior model probabilities were often near 1 when the correct model was chosen and most often multiple models had reasonable support when the incorrect model was chosen. This shows that if there was insufficient information to identify the correct model, the correct model was often still given some posterior support. We find that this method does well even with datasets from 50 outbreaks in households.

In the second case study we found that the data was highly informative for being able to determine the time of symptom onset relative to infectiousness. In the case of the post-symptomatic infection model we selected the correct model every time once 100 households were infected and for the pre-symptomatic infection model we selected the correct model every time once 150 households were infected. In all cases we generally saw an increased ability to select the correct model as more data was obtained. With data on 50 outbreaks and loose priors the Pre model had the lowest proportion of correct selections at 0.7. With data on 150 outbreaks and loose priors the Co model had the lowest proportion of correct selections at 0.9. This shows that symptom onset data from multiple outbreaks is highly informative for choosing the time of symptom onset relative to infectiousness. The results of both studies show that these kinds of data sets are sufficient for discriminating between these models in the early stages of an outbreak of a novel disease, which has important implications for informing public health response [1, 3, 2].

## Acknowledgements

JNW acknowledges the support provided by the Dr Michael and Heather Dunne PhD Scholarship in Applied Mathematics and an Australian Government Research Training Program Stipend Scholarship. JVR acknowledges the support of an Australian Research Council Future Fellowship (FT130100254). AJB was supported by an Australian Research Council Discovery Early Career Researcher Award (DE160100690). All authors also acknowledge support from both the ARC Centre of Excellence for Mathematical and Statistical Frontiers (CoE ACEMS), and the Australian Government NHMRC Centre for Research Excellence in Policy Relevant Infectious diseases Simulation and Mathematical Modelling (CRE PRISM2). This work was supported with supercomputing resources provided by the Phoenix HPC service at the University of Adelaide.

