## Supplementary figures for "Bayesian model discrimination for partially-observed epidemic models"

---

This document provides the supplementary material in support of the paper "Bayesian model discrimination for partially-observed epidemic models". The supplementary material consists of figures displaying some results from the paper.

Bar graphs of 10 randomly-selected sets of posterior model probabilities are given for the SIR, SI(2)R and SI(5)R models in Figures 1-3. Bar graphs of 10 randomly-selected sets of posterior model probabilities based on final outbreak size data from the SIR, SI(2)R and SI(5)R models are given in Figures 4-6. Bar graphs of 10 randomly-selected sets of posterior model probabilities from the Post, Co and Pre SEEIIR models are given in Figures 7-9. Box plots of the differences between the posterior model probability of the true model and the alternative models for the SIR, SI(2)R and SI(5)R model based on final size data are given in Figures 10 and 11.

*Corresponding author

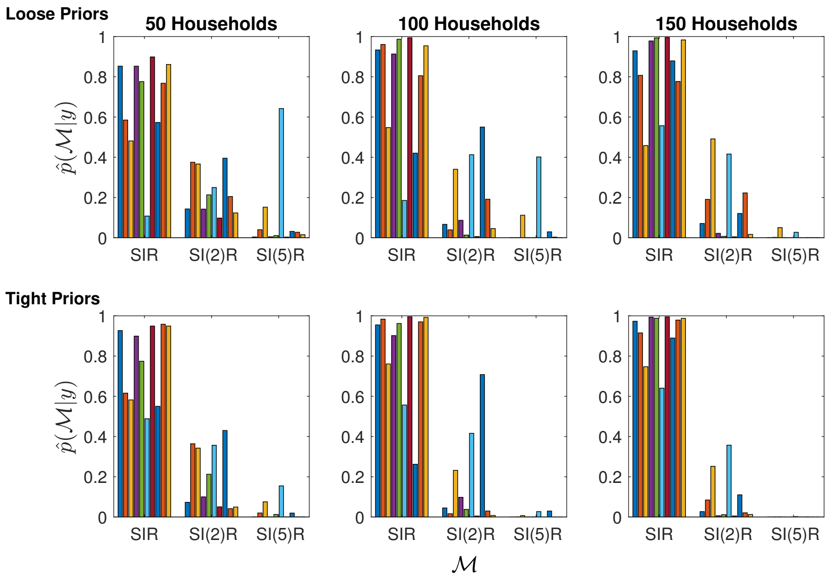


Figure 1: Bar graphs of the posterior model probabilities from 10 data sets generated from the SIR model. The upper and lower panels are results from loose and tight priors respectively. The panels from left to right are results based on 50, 100 and 150 independent outbreaks in households.

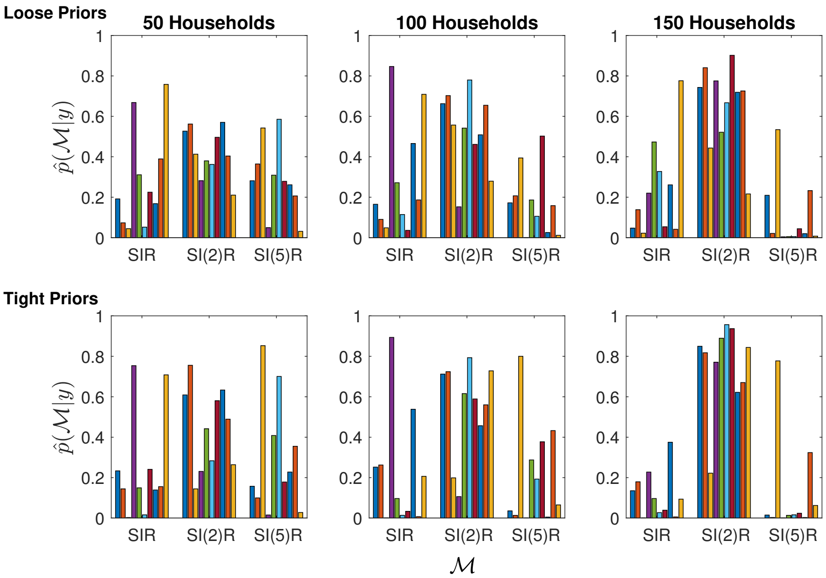


Figure 2: Bar graphs of the posterior model probabilities from 10 data sets generated from the SI(2)R model. The upper and lower panels are results from loose and tight priors respectively. The panels from left to right are results based on 50, 100 and 150 independent outbreaks in households.

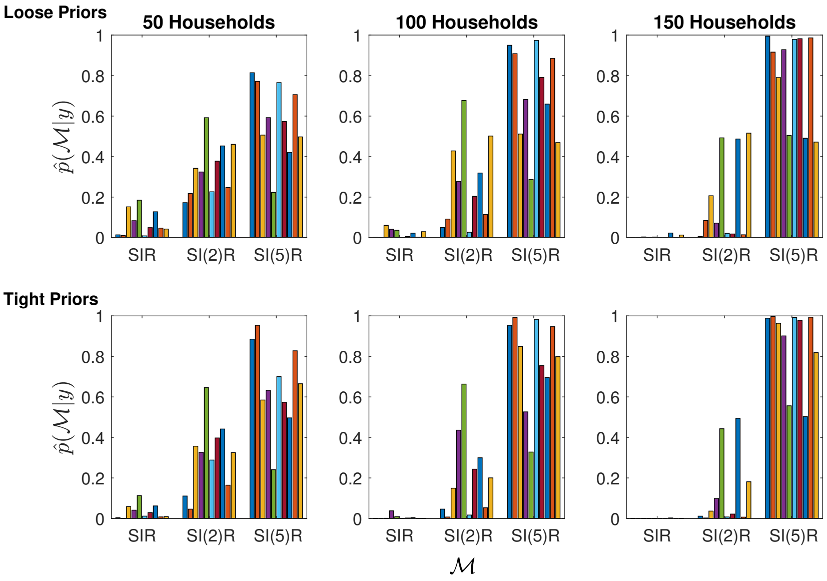


Figure 3: Bar graphs of the posterior model probabilities from 10 data sets generated from the SI(5)R model. The upper and lower panels are results from loose and tight priors respectively. The panels from left to right are results based on 50, 100 and 150 independent outbreaks in households.

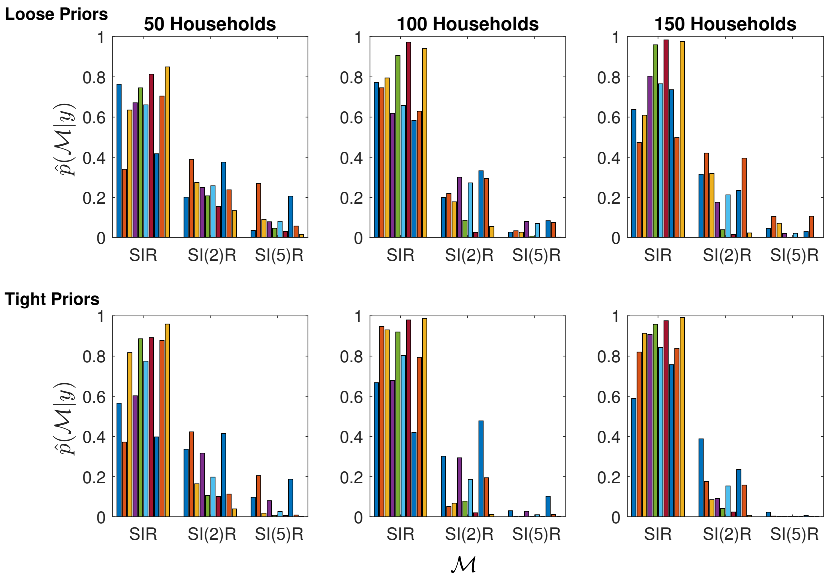


Figure 4: Bar graphs of the posterior model probabilities from 10 data sets of final size data generated from the SIR model. The upper and lower panels are results from loose and tight priors respectively. The panels from left to right are results based on 50, 100 and 150 independent outbreaks in households.

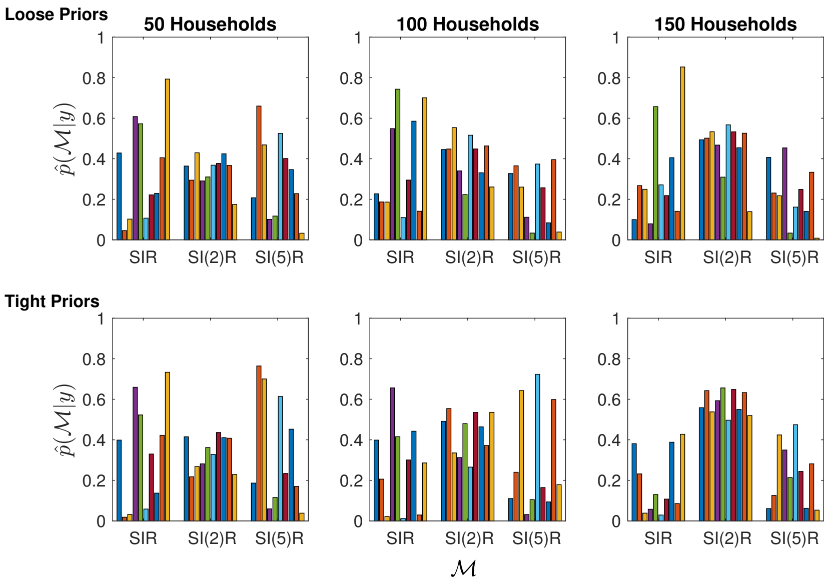


Figure 5: Bar graphs of the posterior model probabilities from 10 data sets of final size data generated from the SI(2)R model. The upper and lower panels are results from loose and tight priors respectively. The panels from left to right are results based on 50, 100 and 150 independent outbreaks in households.

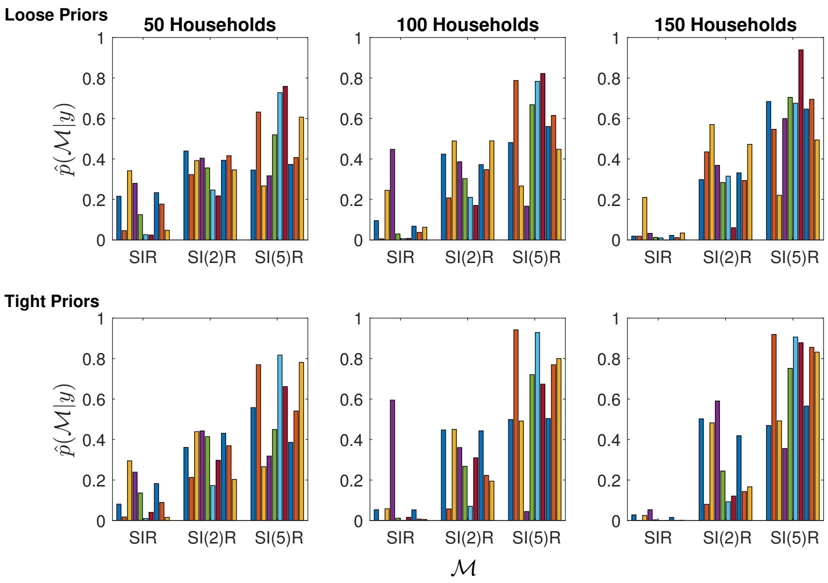


Figure 6: Bar graphs of the posterior model probabilities from 10 data sets of final size data generated from the SI(5)R model. The upper and lower panels are results from loose and tight priors respectively. The panels from left to right are results based on 50, 100 and 150 independent outbreaks in households.

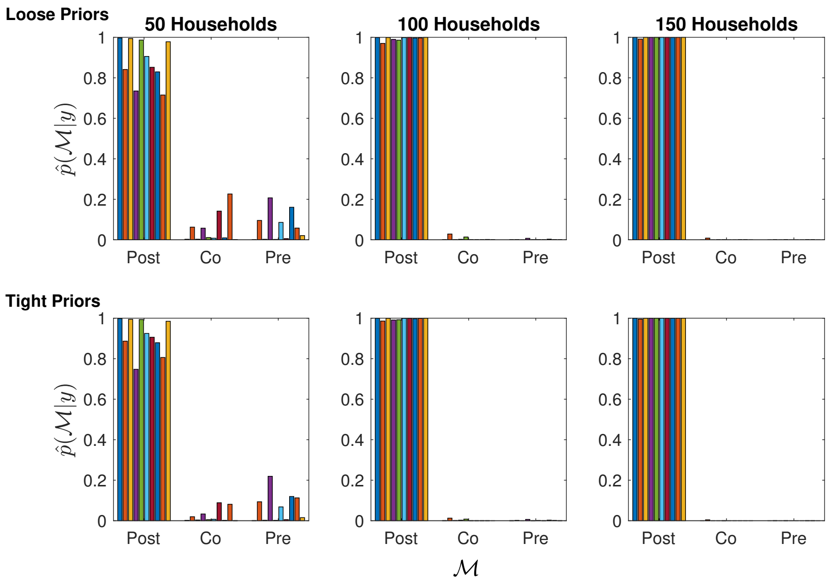


Figure 7: Bar graphs of the posterior model probabilities from 10 data sets generated from the post-symptomatic infection SEEIIR model. The upper and lower panels are results from loose and tight priors respectively. The panels from left to right are results based on 50, 100 and 150 independent outbreaks in households.

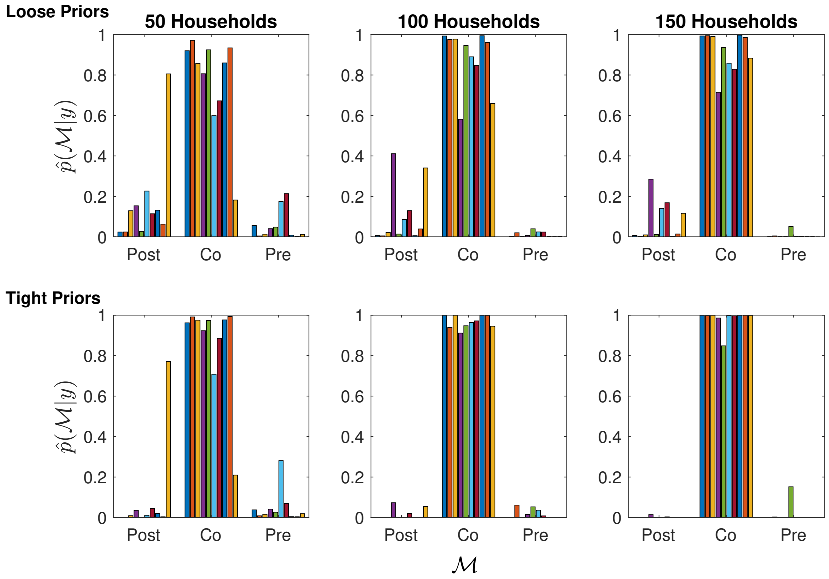


Figure 8: Bar graphs of the posterior model probabilities from 10 data sets generated from the coincidental symptomatic infection SEEIIR model. The upper and lower panels are results from loose and tight priors respectively. The panels from left to right are results based on 50, 100 and 150 independent outbreaks in households.

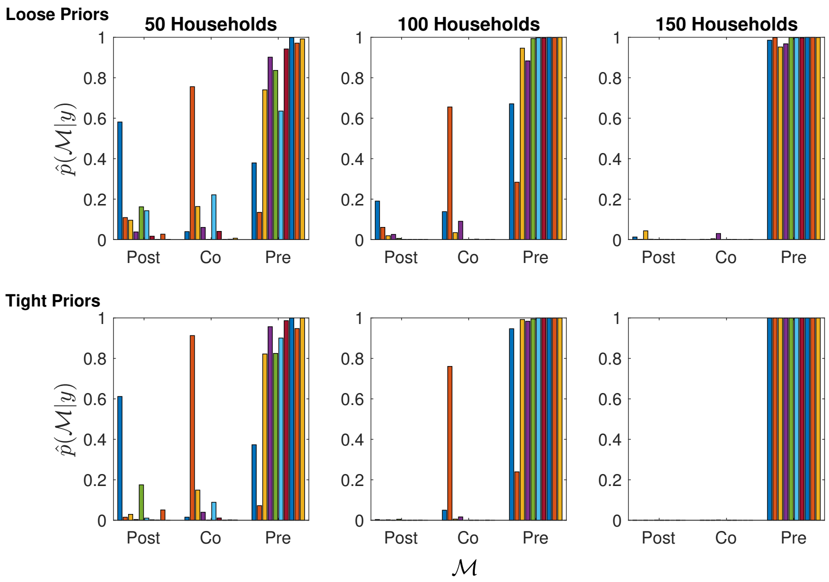


Figure 9: Bar graphs of the posterior model probabilities from 10 data sets generated from the pre-symptomatic infection SEEIIR model. The upper and lower panels are results from loose and tight priors respectively. The panels from left to right are results based on 50, 100 and 150 independent outbreaks in households.

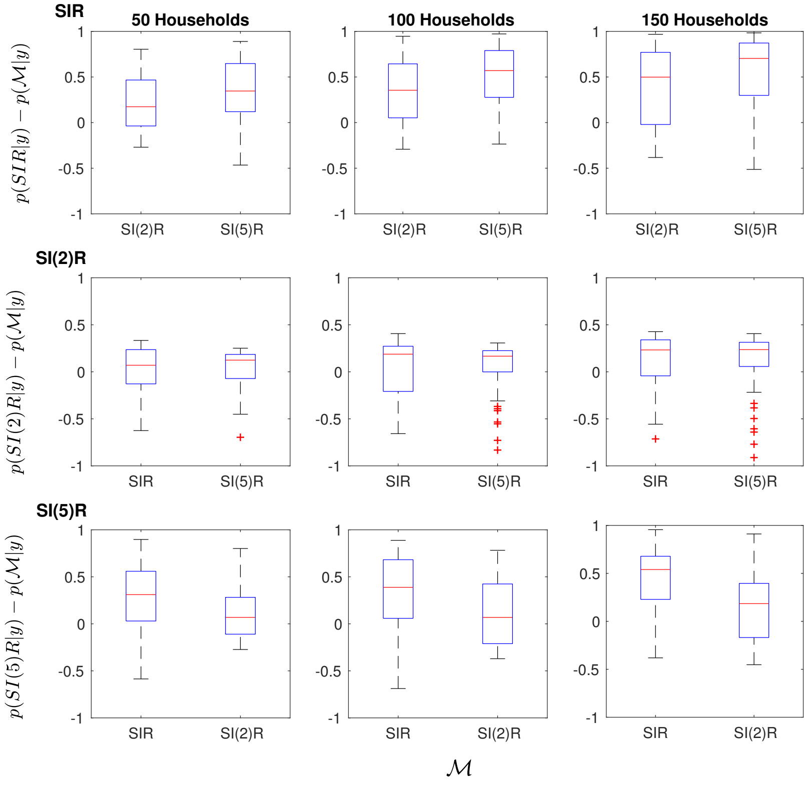


Figure 10: Box plots of the difference in posterior model probability of the true model and the other candidate models for the SI(k)R models with loose priors based on 50 simulated final outbreak size data sets. For example, in the upper left panel, boxes on the left and right of are made using 50 estimates of $p(\text{SIR}|y) - p(\text{SI(2)R}|y)$ and $p(\text{SIR}|y) - p(\text{SI(5)R}|y)$ respectively. Rows from top to bottom show results from data sets generated from the SIR, SI(2)R and SI(5)R models. Columns from left to right represent data sets containing 50, 100 and 150 independent outbreaks in households.

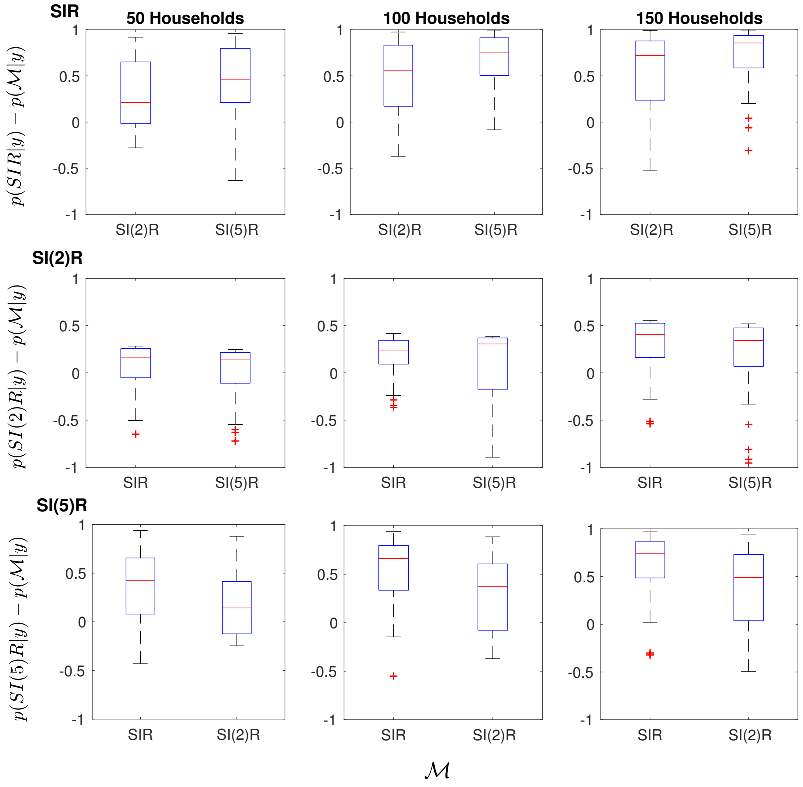


Figure 11: Box plots of the difference in posterior model probability of the true model and the other candidate models for the SI(k)R models with tight priors based on 50 simulated final outbreak size data sets. For example, in the upper left panel, boxes on the left and right of are made using 50 estimates of $p(\text{SIR}|y) - p(\text{SI(2)R}|y)$ and $p(\text{SIR}|y) - p(\text{SI(5)R}|y)$ respectively. Rows from top to bottom show results from data sets generated from the SIR, SI(2)R and SI(5)R models. Columns from left to right represent data sets containing 50, 100 and 150 independent outbreaks in households.
